# Thalamic and cortical signals synergistically represent auditory prediction errors

**DOI:** 10.64898/2026.08.06.743264

**Authors:** Claudia Pascovich, Juho Äijälä, Santiago Castro-Zaballa, Alicia Costa, Alejo Rodriguez-Cattáneo, Pablo Torterolo, Robin A.A. Ince, Tristan A. Bekinschtein, Andres Canales-Johnson

**Affiliations:** Laboratory of Sleep Neurobiology, Department of Physiology, Facultad de Medicina, Universidad de la República, Montevideo, Uruguay; Department of Psychology, University of Cambridge, CB2 3EB Cambridge, United Kingdom; Neuroscience Center, Helsinki Institute of Life Science, University of Helsinki, P.O. Box 3, Fabianinkatu 33, FI-00014 Helsinki, Finland; Institute of Neuroscience and Psychology, University of Glasgow, Scotland G12 8QB, United Kingdom; CINPSI Neurocog, Faculty of Health Sciences, Universidad Católica del Maule, 3460000 Talca, Chile; Department of Clinical Neuroscience, Karolinska Institutet, Stockholm, Sweden

## Abstract

Prediction errors (PEs) are commonly described as cortical signals generated within sensory hierarchies, but whether the thalamus participates in their encoding and transmission remains unclear. We recorded Local Field Potentials (LFP) from the medial and lateral geniculate nuclei and electrocorticography (ECoG) from multiple cortical regions in three awake cats during two auditory prediction tasks. Mutual information (MI) analyses revealed PE encoding in both thalamic and cortical signals. Co-information (co-I) analyses showed off-diagonal temporal synergy between early and later thalamic response components, consistent with an early response inducing a neural state change that shaped the informational content of subsequent activity. Multivariate co-information (MVCo-I) further revealed that thalamic and cortical population activity carried complementary PE information unavailable from either thalamic or cortical areas alone. These synergistic interactions were reliable across animals for violations of structured auditory sequences and weaker for repetition-based deviants. These findings show that auditory PEs are not simply relayed or duplicated across the thalamocortical hierarchy. Instead, they emerge through state-dependent transformations within the thalamus and complementary interactions between thalamic and cortical populations, identifying the thalamus as an active node of context-dependent PE processing.

## INTRODUCTION

The brain continuously compares incoming sensory input with predictions derived from recent experience. When expected and actual events diverge, prediction errors (PEs) provide a signal required to update internal models of the environment. Predictive processing theories propose that these errors are not generated by isolated sensory areas but rather through recurrent interactions between ascending sensory evidence and descending predictions within hierarchical circuits (Aizenbud et al., 2025; Rao and Ballard, 1999; Vinck et al., 2025). Auditory mismatch responses provide a useful model for studying these computations across species: violations of acoustic regularities elicit PE responses along the auditory pathway, including subcortical structures and cortex (Blenkmann et al., 2019; Blume et al., 2026; Canales-Johnson et al., 2021; Gelens et al., 2024; Näätänen et al., 2011; Parras et al., 2017). Indeed, single-neuron recordings have shown that auditory PE signals emerge at subcortical stages of the auditory pathway, with the prevalence and strength of PE coding increasing toward the auditory cortex, supporting a hierarchical organization of deviance detection (Parras et al., 2017). Yet a central question remains unresolved: whether the thalamus merely shares common PE information with the cortex or participates in constructing prediction errors through recurrent thalamocortical interactions.

This question has become especially pressing because recent circuit evidence challenges a feedforward view of how PEs are generated. In the mouse visual cortex, a cooperative thalamocortical disinhibitory circuit is required for generating sensory PE signals, demonstrating that thalamic inputs can actively shape cortical error responses rather than simply relay sensory information (Furutachi et al., 2024). This aligns with anatomical and physiological evidence that corticothalamic pathways are positioned to regulate thalamic gain, selectivity, and context sensitivity (Antunes and Malmierca, 2021). The thalamus is therefore not only a gateway for sensory input, but a recurrent computational hub in which sensory evidence and cortical predictions can interact. What remains unknown is how these interactions are organized in terms of information: whether thalamic and cortical populations carry overlapping information about PEs, or whether their joint activity encodes synergistic information unavailable from either structure alone.

Answering this question requires moving beyond analyzing PE responses as confined to individual neuroanatomical areas, or simple pairwise comparisons between different levels of the hierarchy. Neural responses, as currently understood, are often distributed: task-relevant information can be encoded in patterns spanning multiple cortical sites and regions (Ä ijälä et al., 2026; Allen et al., 2017; Gelens et al., 2024; Musall et al., 2019; Steinmetz et al., 2019). Auditory PE signals likewise occur across subcortical nuclei, the thalamus, auditory cortex, and frontal and association cortices (Carbajal and Malmierca, 2018; Parras et al., 2017; Wacongne et al., 2011). Thus, analyzing each region separately can distinguish whether the region responds to irregularities, but not how the whole system represents the information jointly, and whether different populations encode the same or complementary information. To investigate this distinction, a brain-wide approach that integrates activity across thalamic and cortical sites is required.

Information theory provides a direct way to distinguish these alternatives (Chidichimo et al., 2025; Ince et al., 2017; Rebbin et al., 2025). Mutual information quantifies how strongly neural activity discriminates between expected and unexpected events, whereas co-information separates information interactions into redundancy and synergy. Redundancy indicates that two signals carry overlapping information about the same event; synergy indicates that their joint activity carries information beyond the sum of their individual contributions. This distinction is critical for predictive-processing models. Redundant coding would support the view that prediction errors are broadcast or duplicated across the hierarchy. Synergistic coding, by contrast, would indicate that PEs are constructed through interactions among neural populations, consistent with recurrent accounts of the types and dynamics of information that underlie error representations across the cortical hierarchy. This establishes synergy as a candidate signature of cortical predictive processing. Complementary work in marmoset auditory cortices has shown that event-related potentials (ERPs) and broadband transients preferentially encode synergistic information about PEs, showing that distributed cortico-cortical PEs can be highly synergistic (Gelens et al., 2024). However, these studies leave unresolved whether synergistic PE coding is specifically cortical or extends into thalamocortical circuits, where predictive signals may first be shaped.

Here, we tested whether auditory PEs are represented through synergistic thalamocortical interactions in awake cats. We used two complementary paradigms because they instantiate distinct forms of auditory expectation. In the Roving Odd-ball task, predictions emerge gradually from stimulus repetition, and violations signal a change in recent sensory statistics (Gelens et al., 2024; Komatsu et al., 2015). By contrast, the Local Effect of the Local–Global paradigm depends on the position of a sound within a structured sequence and therefore probes violations of an immediate temporal rule (Bekinschtein et al., 2009; Gelens et al., 2024). Comparing these paradigms allowed us to determine whether thalamic and thalamocortical synergy reflects a general property of deviance processing or varies with the structure and timescale of the underlying prediction. Using recordings from the medial and lateral geniculate nuclei, auditory cortex, prefrontal cortex, and additional cortical sites, we first quantified PE information with mutual information and then used co-information and multivariate co-information to distinguish redundant from synergistic representations within thalamic signals and across thalamocortical populations. We hypothesized that, if the thalamus participates in recurrent predictive inference rather than simply relaying cortical or sensory signals, thalamic activity should encode PE information synergistically across time, reflecting the dynamic influence of recurrent inputs from the cortex and other regions. Furthermore, the thalamus should contribute complementary information to cortical PE representations, as it interacts with cortical PE-encoding. This design, therefore, tests both the presence and the contextual dependence of synergistic thalamocortical coding.

## RESULTS

### Mutual information reveals PE responses within thalamic and cortical areas

To characterize PE dynamics across the thalamocortical system, we recorded electrocorticography (ECoG) signals from the auditory, prefrontal, motor, parietal, and visual cortices, as well as Local Field Potential (LFP) signals from the medial and lateral geniculate nuclei, in three cats. From the ECoG signals, we then calculated ERPs during two auditory paradigms: a Roving Oddball task and the Local-Global task, with a focus on the Local Effect. The auditory task designs, stimuli, and recording techniques were preregistered (https://osf.io/tfjwg). An illustration of the experimental design is depicted in Figure 1.

**Figure 1:**
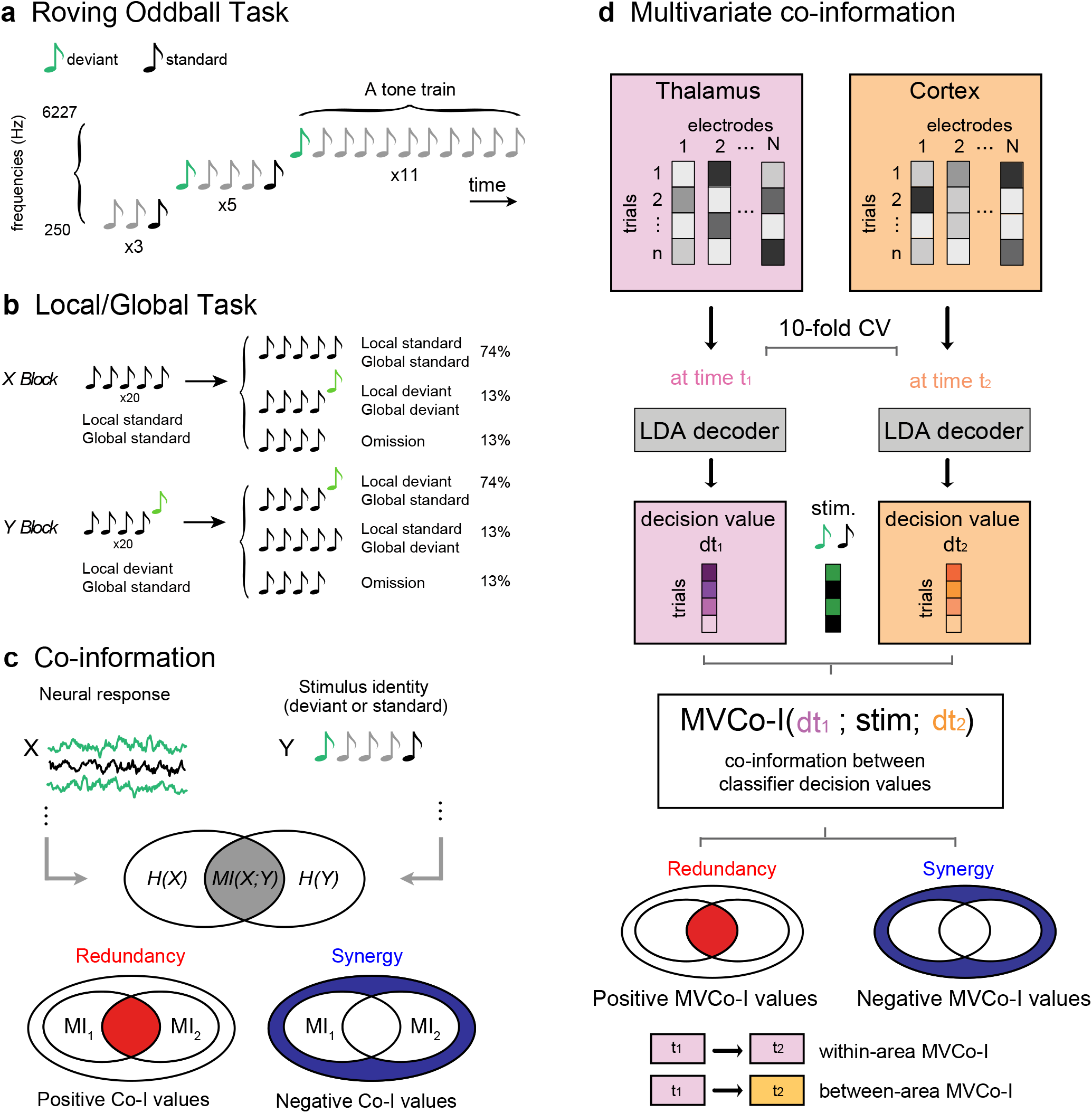
Schematic illustrating the experimental design and presentation of the stimuli. **(A)** Roving task. Twenty different unique tones are presented in trains of 3, 5, or 11 identical stimuli. Either of the two later trains consisted of different tones. In this way, while the adjacent standard (represented in blue) and offset tones (represented in magenta) shifted in frequency due to the transition between the trains, the two expectation conditions physically coincided, since the first and last tone from the same train are treated as deviated and standard tones in the analysis of pairs of adjacent stimuli. **(B)** Local-Global task. In each Trial, 5 complex sounds of 50 ms duration were presented. Four different types of sound series were used; in the first 2 the same 5 sounds were used (EEEEE or OOOOO), and the second series of sounds was EEEEO, OOOOE in the case of the deviant, or EEEE or OOOO for the omission. **(C)** Co-information analysis. Mutual information quantified the information carried by each neural signal about stimulus identity. Positive co-information indicates redundant information shared by two signals, whereas negative co-information indicates synergistic information available only from their joint activity. **(D)** Multivariate co-information analysis. Linear discriminant classifiers were trained with 10-fold cross-validation on multichannel activity from thalamic or cortical electrode populations. Out-of-sample classifier decision values were used to quantify the mutual information each population carries about stimulus identity. MVCo-I was then computed between decision values obtained at different time points, either within the same multivariate population or between thalamic and cortical populations. Positive MVCo-I indicates information redundantly accessible from both multivariate representations, whereas negative MVCo-I indicates complementary information that becomes available only when the two representations are considered jointly.

To examine how PE was distributed across cortical and thalamic regions, we quantified PE at each electrode by comparing responses to deviant and standard tones. For each electrode, we calculated MI to assess the dependence between tone category (i.e., standard vs. deviant) and the corresponding neural signal across trials. In information theory, MI is a statistical measure of the strength of dependence between two variables and can also be interpreted as an effect size (measured in bits) for a test of statistical independence (Ince et al., 2017). Accordingly, for each electrode and time point, we extracted ECoG signals from standard and deviant trials and used MI to quantify the effect size of the difference between them.

A well-studied ERP marker of auditory PE is the mismatch negativity (MMN), an ERP component in the auditory cortex of the cat that peaks around 30-70 ms after the onset of an infrequent acoustic stimulus and is followed by a large positivity lasting from 60 to 120 ms (Csépe et al., 1987). Within the framework of predictive processing, MMNs can be interpreted as the difference between the brain’s prediction (i.e., the internal model) and the observed sensory signal (Chennu et al., 2013; Friston, 2005; Garrido et al., 2008, 2009b; Winkler et al., 2009), i.e., a PE response. MI was computed for the PE signal recorded in each cortical and thalamic electrode. The ERPS and MI dynamics over time in the lateral or medial geniculate nuclei and in the auditory and prefrontal cortices are shown in Figure 2. For both paradigms, the Roving Oddball Task and the Local-Global Task, ERP signals showed PE effects across multiple cortical regions not necessarily restricted to canonical auditory areas (Figure 2).

**Figure 2:**
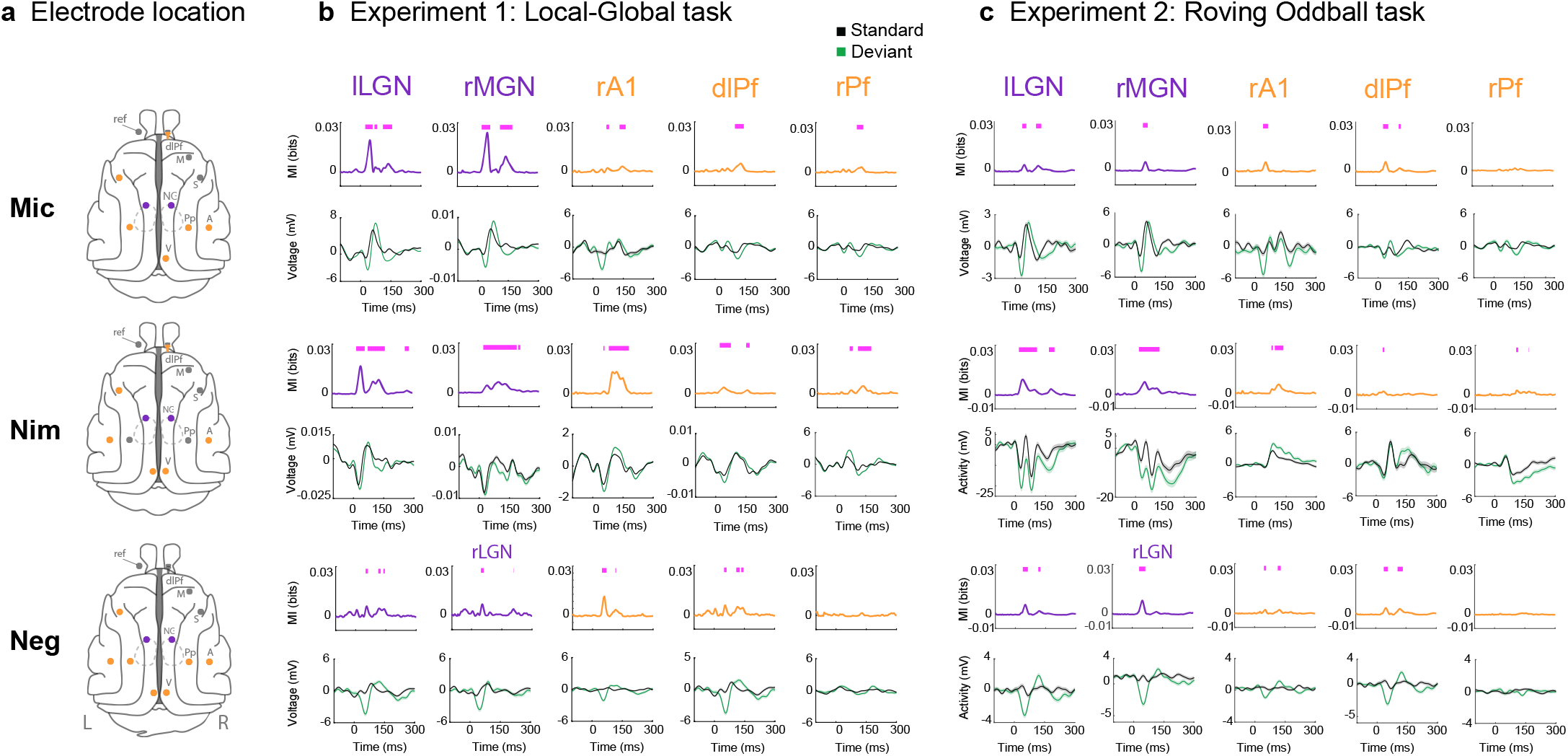
ERP markers of PE across the cat brain. **(A)** Electrode locations for cat Mic, Nim, and Neg in Experiments 1 and 2. Electrodes showing significant PE effect after computing MI between standard and deviant trials for the ERP markers of auditory PE are shown as red circles in the respective cat brain illustrations. Experiment 1: Global–Local task. PE signals (MI and ERPs) in the auditory network, including the medial and lateral geniculate nuclei (MGN and LGN), primary auditory cortex (A1), and rostral and dorsolateral prefrontal cortex (rPf and dlPf). MI values in bits (effect size of the difference) are shown first, followed by the corresponding ERP responses to deviant tones (green) and standard tones (black) for the same locations. **(C)** Experiment 2: Roving Oddball task. MI and ERP responses are shown for the same locations and using the same conventions as in **(B).** The letters r or l indicate right or left, respectively. Significant time points after the permutation test are shown as fuchsia bars over the MI plots. Significant time windows shown were computed using nonparametric permutation testing controlling FWER < 0.05 over all time points in the trial.

### Co-information reveals redundant and synergistic PE representations within thalamic signals

Having established that the medial and lateral geniculate nuclei encoded PE information, we next examined the temporal information structure of these thalamic responses using co-information analysis (co-I) (Ince et al., 2017). When computed over time, co-I distinguishes information about stimulus category that is redundantly represented across two time points from information that becomes available only when those time points are considered jointly. Positive co-I, therefore, indicates temporally redundant PE information, whereas negative co-I indicates temporal synergy (Figure 1c). The analysis was performed within individual thalamic ERP signals and was restricted to electrodes exhibiting significant mutual information effects (Figure 2).

For the Local Effect, MI in the LGN and MGN showed a prominent early component around 50 ms, followed by additional components around 90–100 ms and 110–140 ms after deviant onset (Figure 3a,b). These temporally distinct PE components were reflected in positive co-I concentrated along and near the diagonal, indicating that overlapping PE information was expressed across neighboring response times. In addition, both thalamic nuclei exhibited off-diagonal negative co-I, particularly between earlier and later portions of the response. These synergistic interactions can be interpreted as an early thalamic response initiating a change in neural state that shaped how PE information was represented at later time points, such that the evolving response jointly conveyed information unavailable from either temporal component alone, and it’s consistent with findings observed within cortical PE signals (Gelens et al., 2024).

**Figure 3:**
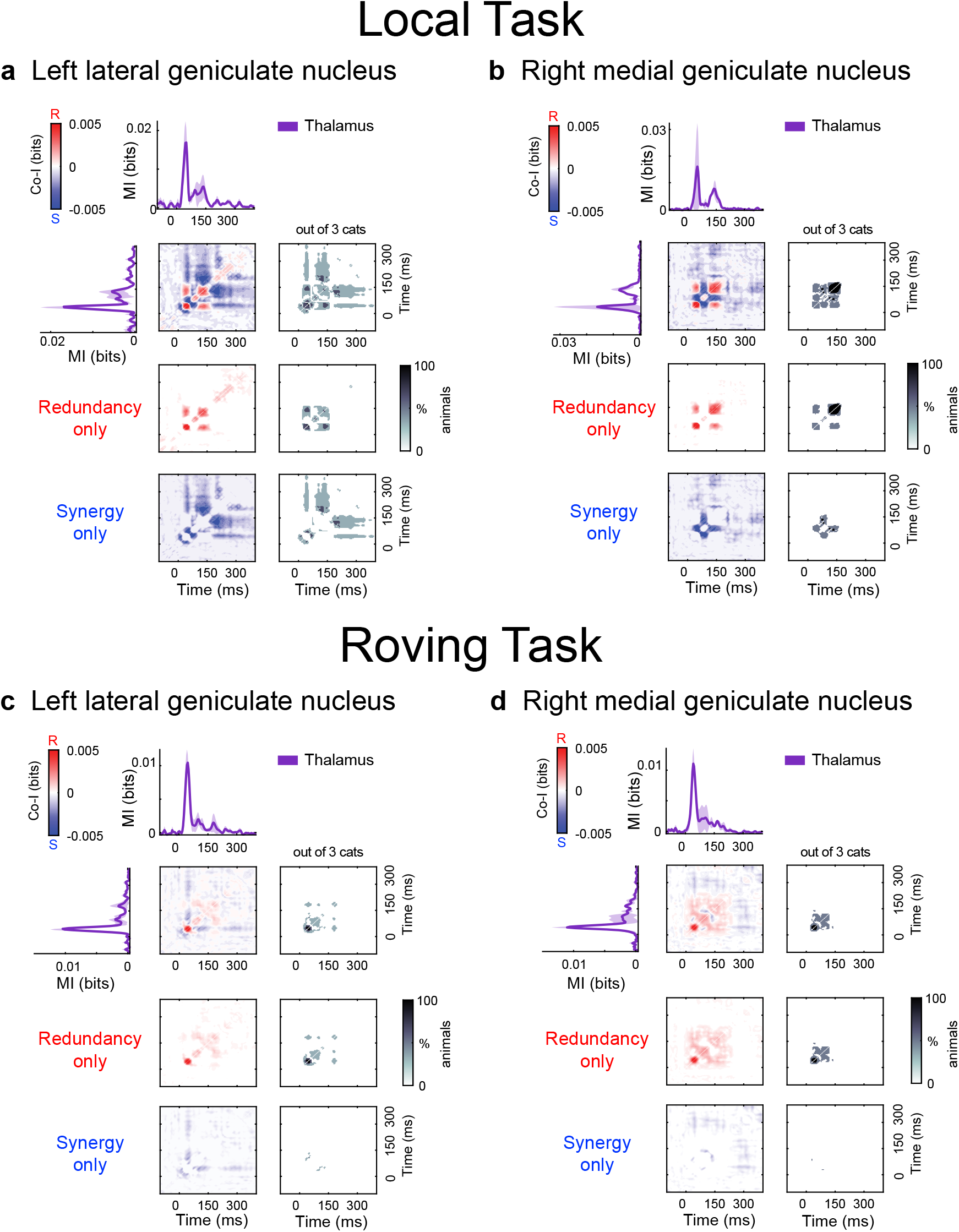
Synergy and redundancy within ERP signals in the thalamic electrodes for the Local Effect of the Local-Global, and Roving Oddball Tasks. **(A)** Co-I revealed synergistic and redundant patterns within ERP signals in the lateral and medial geniculate nuclei for Local (Panels **a** and **b**), and for the Roving Task (Panels **c** and **d**). MI (solid traces) between standard and deviant trials for each electrode (one per cat) averaged across the three cats. co-I was computed within the corresponding signal (ERP) across time points between -100 and 350 ms after tone presentation. The average of the corresponding electrodes across cats is shown for the complete co-I chart (red and blue plots), for positive co-I values (redundancy only; red panel), and for negative co-I values (synergy only; blue plot). The grey-scale plots show the portion of cats exhibiting significant co-I differences in the single-electrode analysis.

The Roving Oddball task also elicited PE information in the lateral and medial geniculate nuclei (Figure 3c,d). Although multiple components were visible in individual animals, the across-animal average was dominated by an early MI peak around 50 ms. Co-I nevertheless revealed both diagonal redundancy and off-diagonal synergy within the thalamic responses. The synergistic effects were concentrated primarily within the early post-stimulus period and were weaker and less consistent across animals than those observed for the Local Effect. Together, these results show that thalamic PE responses are not composed solely of independent or repeated information over time. Instead, their information structure includes synergistic interactions that enable distinct phases of the thalamic response to jointly encode auditory violations.

### Multivariate co-I reveals temporally distributed PE representations across thalamic and cortical areas

After characterizing the information structure of individual thalamic signals, we next used multivariate co-information (MVCo-I) (Gelens et al., 2024) to examine how distributed PE information was represented across the combined population activity of the thalamus and all recorded cortical electrodes (see Methods). At each time point, multivariate classifiers were trained to discriminate standard from deviant trials using the joint activity of all thalamic and cortical electrodes. Mutual information between the cross-validated classifier decision values and the stimulus category confirmed that this distributed thalamic-cortical activity encoded PE information in both paradigms, although the information was stronger and more temporally sustained in the Local Effect than in the Roving Oddball task.

For the Local Effect, temporal MVCo-I revealed both redundant and synergistic components within the distributed thalamic–cortical representation (Figure 4a). Redundant information was concentrated along the diagonal and extended between neighboring time points, indicating that overlapping PE information was preserved across successive time points. Additional off-diagonal redundancy showed that common PE information was also accessible from multivariate patterns expressed at more temporally separated stages. In parallel, widespread off-diagonal negative MVCo-I revealed synergy between earlier and later population representations. Thus, combining multivariate activity across different time points provided additional information about the violation that was unavailable from either representation considered in isolation. The individual animal charts (Supplementary Figures S1–S3) showed significant synergy in all three cats (3/3) and significant redundancy in two of the three cats (2/3). The Roving Oddball task yielded a weaker, less consistent multivariate information structure (Supplementary Figure S4). Multivariate MI was significant in two of the three cats (2/3), but neither significant redundancy nor significant synergy was detected in any cat (0/3 for both redundancy and synergy). These findings indicate that, although PE information could be decoded from the combined thalamo–cortical signals during the Roving Oddball task, there was no significant evidence of temporal redundancy or synergy, i.e., distributed coding of PE information.

**Figure 4:**
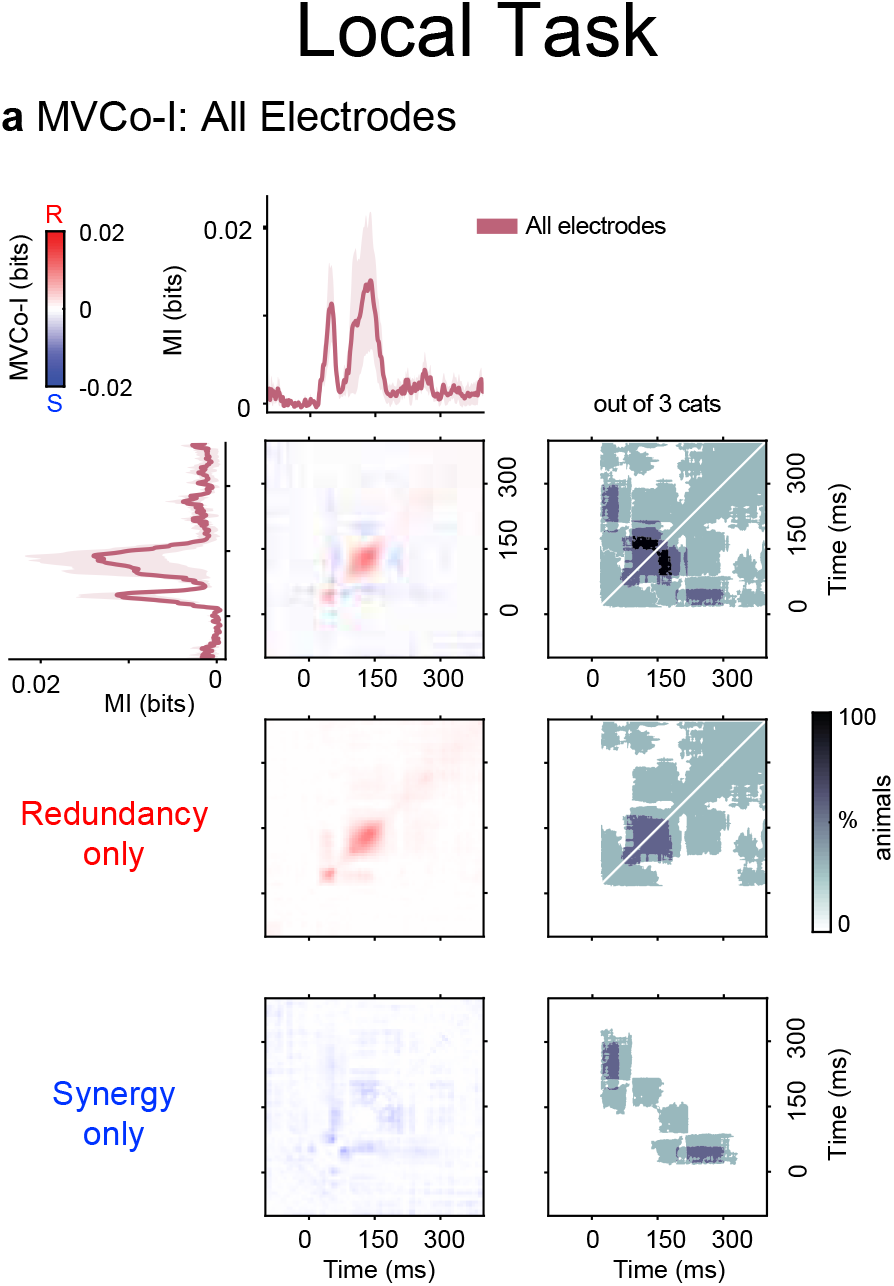
Multivariate redundancy and synergy within distributed thalamic and cortical PE representations during the Local task. MVCo-I was used to characterize the temporal information structure of distributed prediction-error representations across all thalamic and cortical electrodes during the local task. At each time point, a multivariate classifier was trained using the joint activity of all thalamic and cortical electrodes to discriminate deviant from standard trials. The solid magenta trace shows the mutual information between the cross-validated classifier decision values and stimulus category, averaged across the three cats. Temporal MVCo-I was calculated between classifier decision values obtained at every pair of time points from -100 to 350 ms relative to tone onset. Positive MVCo-I values indicate redundant PE information expressed across time, whereas negative values indicate synergistic information available only when the multivariate representations at two time points are considered jointly. The central matrix shows the complete MVCo-I values, with redundancy in red and synergy in blue. The lower matrices separately display only positive values (redundancy) and only negative values (synergy). Grey-scale matrices indicate the proportion of cats showing statistically significant MVCo-I at each pair of time points. Statistical significance was assessed using nonparametric permutation testing with family-wise error-rate correction at *p* < 0.05.

### Multivariate co-I reveals synergistic PE representations between thalamic and cortical areas

We subsequently tested whether PE information was redundantly shared or synergistically integrated between the thalamus and the network formed by all recorded cortical electrodes. Separate multivariate classifiers were trained on the thalamic and all-cortical electrode populations, and MVCo-I was calculated between their cross-validated decision values at every pair of time points (Figure 1d). For the local effect of the LocalGlobal task (Figure 5), both the thalamic classifier and the classifier trained on all cortical electrodes contained information distinguishing standard from deviant trials, but their temporal profiles differed. Thalamic PE information was strongest relatively early in the post-stimulus period, whereas the cortical representation derived from all recorded cortical electrodes was more broadly distributed across subsequent time points. Between-region MVCo-I was predominantly synergistic and off-diagonal, demonstrating that thalamic activity at one stage of processing and cortical activity at another stage made complementary contributions to the representation of the local PE-encoding. Their joint multivariate state, therefore, carried more PE information than could be recovered from either regional representation alone. Positive MVCo-I was comparatively sparse, indicating that only a limited portion of PE information was redundantly accessible from both the thalamic and cortical populations. By contrast, synergistic interactions extended across a broad range of thalamic–cortical time-point combinations. The individual animal MVco-I maps (Supplementary Figures S5–S7) showed significant synergy in all three cats (3/3), whereas significant redundancy was present in two of the three cats (2/3). These results therefore show that the encoding of local violations of regularity was not simply duplicated across thalamic and cortical networks. Instead, it emerged primarily through temporally distributed, complementary interactions between their population-level representations, consistent with the participation of recurrent thalamocortical computations in constructing auditory prediction errors.

**Figure 5:**
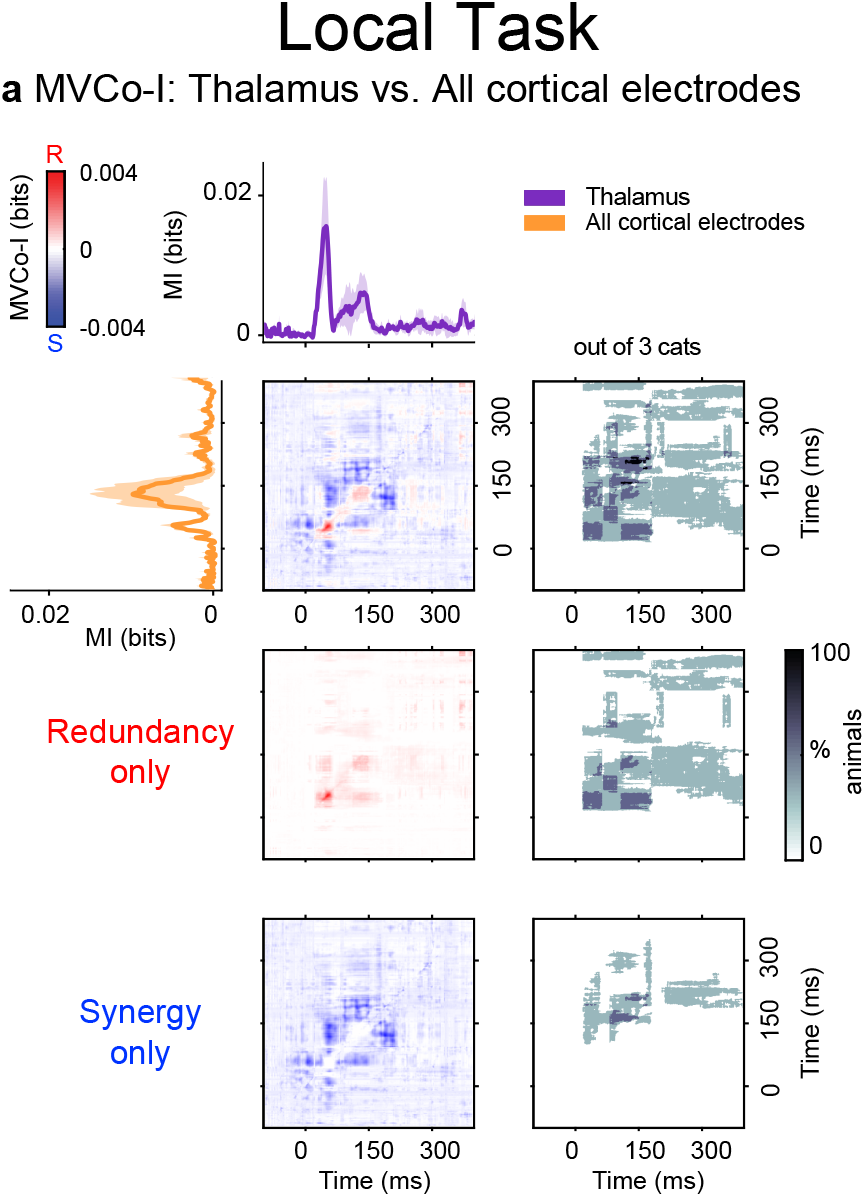
Multivariate redundancy and synergy between thalamic and cortical PE representations during the Local task. MVCo-I quantified the informational relationship between multivariate prediction-error representations in the thalamus and those across all cortical electrodes during the local task. Separate multivariate classifiers were trained at each time point using either thalamic electrodes or all cortical electrodes to discriminate deviant from standard trials. The black and orange traces show the mutual information between the cross-validated classifier decision values and stimulus category for the thalamic and cortical models, respectively, averaged across the three cats. Cross-regional MVCo-I was calculated between thalamic decision values at each time point and cortical decision values at every other time point from -100 to 350 ms relative to tone onset. Positive MVCo-I values indicate PE information redundantly accessible from both regional representations, whereas negative values indicate synergistic information available only from their joint multivariate state. The central matrix shows the complete MVCo-I values, with redundancy in red and synergy in blue. The lower matrices separately display only positive values (redundancy) and only negative values (synergy). Grey-scale matrices indicate the proportion of cats showing statistically significant MVCo-I at each thalamic–cortical time-point combination. Statistical significance was assessed using nonparametric permutation testing with family-wise error-rate correction at *p* < 0.05.

The Roving Oddball task produced a weaker and less consistent between-region information structure (Supplementary Figure S8). Although the MVCo-I contained localized redundant and synergistic components, significant synergy was not detected in any cat (0/3), while significant redundancy was detected in only one cat (1/3). Thus, the complementary thalamic–cortical interactions observed during the Local Effect were not consistently expressed during purely repetition-based deviance processing.

## DISCUSSION

In this study, we investigated how PE signals are distributed and integrated across thalamocortical signals during wakefulness in cats. By combining ERPs with information-theoretic analyses, we reveal that auditory PEs are encoded by synergistic and redundant dynamics spanning the thalamus and multiple cortical regions. Our findings demonstrate that PE processing in the thalamocortical system is informationally synergistic rather than merely distributed, supporting a model in which recurrent, hierarchical interactions give rise to adaptive auditory predictions.

### Synergistic encoding of prediction errors across thalamocortical hierarchies

In the case of the information observed across time within thalamic signals, the medial and lateral geniculate nuclei (MGN, LGN) exhibited clear synergistic co-I patterns, suggesting that thalamic structures actively participate in constructing error representations rather than passively relaying sensory input. The off-diagonal synergy between early and later thalamic responses suggests that PE processing involved a sequence of state-dependent transformations rather than a succession of independent response components. In this interpretation, the early response to the violation changes the state of the thalamic network, thereby altering the informational meaning or readout of subsequent activity. Consequently, later thalamic activity becomes informative not only through its instantaneous amplitude, but through its relationship to the preceding response state. This interpretation follows previous work showing that temporal synergy can index a neural state change, whereby activity at an earlier processing stage modifies how stimulus information is represented or decoded at a later stage (Gelens et al., 2024).

This early emergence of synergy is consistent with anatomical and physiological data showing dense reciprocal corticothalamic projections from layers 5 and 6 of auditory and association cortices (Malmierca et al., 2015). These feedback pathways are thought to convey predictive signals that shape thalamic activity, enabling fine-grained comparison between expected and incoming sensory information (Kommajosyula et al., 2021; Lesicko et al., 2022). The synergistic encoding observed here may thus reflect the iterative updating of generative models, whereby thalamic and cortical signals jointly encode unexpected deviations from predicted auditory sequences. PEinformation was simultaneously present and decodable within both thalamic and cortical sites, from an overlapping window of approximately 50 ms to 125 ms after stimulus presentation, linking the early thalamic peak response at approximately 50 ms to the later cortical peak response at 125 ms. Noteworthily, the cross-temporal interaction between these two peaks was primarily synergistic, indicating that the information from the thalamus was not simply duplicated in the cortical response. Rather, the two distinct regions provided complementary information across different processing stages, consistent with recurrent thalamocortical construction of prediction errors.

On the other hand, the presence of synergistic information between thalamic and cortical signals indicates that PE representations arise from the joint computation across multiple brain regions, in which the combined information exceeds the sum of individual contributions. This finding challenges a purely feedforward account of PE generation and instead supports recurrent predictive processing within the thalamocortical hierarchy (Gelens et al., 2024; Shipp, 2016; Usrey and Sherman, 2023). Our results extend recent studies showing distributed synergy in cortical PE signals (Gelens et al., 2024), suggesting that synergistic information processing is a conserved computational motif of hierarchical sensory systems. Importantly, we demonstrate that this motif is already present at the thalamic level, underscoring the thalamus’s active role in the generation and refinement of PEs (e.g., via a specific thalamocortical disinhibitory circuit) (Furutachi et al., 2024).

### Context-dependent modulation of synergy and redundancy

The balance between synergy and redundancy varied across experimental paradigms and brain regions, indicating that the informational architecture of PE processing is dynamically modulated by task context. In Oddball paradigms, context is defined by the learned statistical regularities of the stimulus environment; therefore, using different paradigms alters the expectations the brain forms and how violations of those expectations are detected (Chennu et al., 2013; Friston, 2005; Garrido et al., 2009a; Winkler et al., 2009). Although both the Roving Oddball and Local-Global paradigms elicit auditory prediction errors, they impose qualitatively different forms of contextual structure. In the Roving paradigm, context is defined by the accumulation of stimulus-specific regularities across repetitions, such that expectations emerge gradually from recent sensory history and prediction errors primarily signal changes in the underlying stimulus statistics (Balaguer et al., 2011; Garrido et al., 2009a). In contrast, the Local Effect of the Local-Global paradigm defines context through a short-term sequential structure established within each trial, thereby generating position-dependent expectations over the course of a single sound sequence (Bekinschtein et al., 2009; Chennu et al., 2013). As a result, prediction errors in the Local Effect reflect violations of immediate temporal structure rather than longer-term statistical regularities.

In the Local Effect of the Local–Global paradigm, we observed robust synergistic interactions within distributed thalamocortical representations and between thalamic and cortical populations. These findings indicate that, even over the short timescale of local auditory expectations, prediction errors are represented through complementary information distributed across the thalamocortical system rather than within isolated regions. Violations of immediate sound-sequence structure elicited joint informational patterns that were not recoverable from either thalamic or cortical population activity alone, consistent with the idea that recurrent thalamocortical interactions contribute to constructing fast sensory prediction errors (Carbajal and Malmierca, 2018; Chennu et al., 2016).

In contrast, the Roving Oddball paradigm (Supplementary Figures S4 and S8), which relies on repetition-based expectations, produced weaker and less consistent multivariate interactions, with no reliable evidence of thalamocortical synergy across animals. This suggests that distributed synergistic coding is not an obligatory feature of auditory prediction errors but depends on the structure of the predictive context. When expectations are generated by more structured sequential regularities, complementary interactions across thalamic and cortical populations appear to play a greater role than when expectations arise primarily through stimulus repetition. Such context-dependent differences are consistent with predictive processing accounts in which the complexity and precision of environmental regularities influence how prediction errors are represented across hierarchical circuits (Friston, 2005; Keller and Mrsic-Flogel, 2018; Parr et al., 2018; Rao and Ballard, 1999; Vinck et al., 2025).

Together, these results show that synergistic thalamocortical PE representations were more prominent during the Local Effect than during the Roving Oddball task. This task dependence parallels the findings of Äijälä et al. (2026), in which diverting attention away from the oddball task reduced thalamic synergy and flattened cortical PE-learning trajectories. Although the manipulations differ, both studies suggest that synergistic PE coding is not an invariant feature of auditory prediction-error processing but depends on the computational context in which predictions are generated.

### Functional implications for predictive computation

The synergistic relationships observed across thalamic and cortical regions imply that PEs are not represented locally, but rather emerge from distributed, recurrent computations involving multiple nodes of the auditory hierarchy (Äijälä et al., 2026; Gelens et al., 2024). Based on neurocomputational work with brain-constrained neural networks, synergy has been proposed to act as an information-theoretic marker of recurrent processing in auditory oddball tasks. (Gelens et al., 2024), indicating when the joint activity of two regions conveys novel information unavailable to either region alone. This complements the traditional view of mismatch responses as localized error signals by positioning PE as a network property arising from the interaction between sensory and associative systems.

The thalamocortical system, therefore, functions as a cooperative computational circuit: cortical feedback conveys predictions that modulate thalamic gain and selectivity, while thalamic output refines cortical estimates by providing context-sensitive error updates (Furutachi et al., 2024). The synergy observed here between thalamic and cortical signals may, then, enable the brain to efficiently encode deviations in a dynamic environment, minimizing redundancy and optimizing representational capacity. Notably, synergistic PE coding was not restricted to auditory areas but was also present in the cortical-wide analysis, suggesting that different sensory regions, as well as multimodal and associative regions, contribute to the higherorder interpretation of unexpected events.

### Comparative and translational significance

The demonstration of synergistic PE processing in a nonhuman species provides crucial comparative evidence for the evolutionary conservation of predictive coding principles. The thalamocortical dynamics observed here in cats closely mirror findings in marmoset and human data (Äijälä et al., 2026; Gelens et al., 2024; Roberts et al., 2026), indicating that synergy-based integration may constitute a fundamental mode of information representation for hierarchical perception across mammals. Moreover, because mismatch negativity (MMN) and Local Effect are widely used biomarkers of sensory prediction and cognitive function, understanding their informational structure at the thalamocortical level could inform translational research on consciousness and neuropsychiatric disorders, in which predictive processing is disrupted.

### Future directions

While work combining intracranial electrophysiology and computational modeling has linked the synergy to recurrent processing across different levels of the hierarchy (Gelens et al., 2024), and the balance of redundancy and synergy to prediction error learning (Äijälä et al., 2026), further work on biological neural systems is needed to establish causal mechanisms that can give rise to synergistic encoding in brain networks. For example, future work that combines the current framework with directed connectivity measures or causal perturbations (e.g., optogenetic silencing of corticothalamic feedback) could clarify whether the observed synergy reflects topdown predictions, bottom-up error signaling, or both. Additionally, expanding recordings to include subcortical nuclei such as the inferior colliculus would help determine whether synergistic encoding originates subcortically or is established through cortical feedback loops.

Another promising direction is to investigate how behavioral or cognitive state modulates informational interactions. Prior research shows that wakefulness, sleep, anesthesia, and attention differentially affect the propagation of PEs (Äijälä et al., 2026; Bekinschtein et al., 2009; Canales-Johnson et al., 2021). Applying the present framework across states could reveal how synergy relates to levels of consciousness and the integration of sensory evidence into perceptual awareness.

## Conclusion

Together, our findings provide direct evidence that auditory PEs are represented synergistically across thalamocortical signals in the awake cat brain. These synergistic interactions reveal that the information conveyed by combined thalamic and cortical activity exceeds the sum of their individual contributions, pointing to a distributed, recurrent, and integrative mechanism for predictive processing. We propose that such synergistic computation constitutes a fundamental principle of thalamocortical organization, supporting adaptive prediction and efficient information processing across the sensory hierarchy.

## Authorship contributions

Conceptualization: C.P., T.B., and A.C.-J. Data analysis: P. and J.Ä . Cat experiments: C.P., S.C-S., A.C., A.R-C, and P.T. Software and methods: R.I., J.Ä ., and A.C.-J. Visualization: C.P., J.Ä ., and A.C-J. Writing original draft: C.P., J.Ä ., and A.C.-J. Editing: T.B., P.T., S.C-S, A.C., A.R-C and R.I.. Supervision: T.B., A.C.-J. and P.T.. Funding acquisition: P.T., A.C., T.B., and A.C-J.

## METHODS

### Animals

The experiments in cats were carried out in the Sleep Neurobiology Laboratory of the Faculty of Medicine, in Uruguay. Three adult cats were used in this study. The animals were obtained from the Institutional Animal Care Facility of the School of Medicine, Universidad de la República, Uruguay, and were determined to be in good health. All experimental procedures were conducted in accordance with the Guide for the Care and Use of Laboratory Animals (8th edition, National Academy Press, Washington, DC, 2011) and were approved by the Institutional and National Animal Care Commissions of the Universidad de la República, Uruguay (Exp. N° 070151-000035-23). Institutional Ethics Committee: https://www.chea.edu.uy/node/29. Adequate measures were taken to minimize pain, discomfort, or stress to the animals. In addition, all efforts were made to use the minimum number of animals necessary to produce reliable scientific data.

### Surgical procedure

The anesthetic procedures were performed with the assistance of a veterinary anesthetist. Prior to surgery, after establishing venous access, Meloxicam (0.2 mg/kg, i.v.) was administered, as well as Dexmedetomidine (3 µg/kg to 5 µg/kg, i.m.), Ketamine (2 mg/kg, i.m.), and Ceftriaxone (30 mg/kg, i.m.). Anesthesia was induced with Propofol via an infusion pump until intubation (5 mg/kg, i.v.). Intubation was performed using a veterinary laryngoscope No. 2. Anesthesia was maintained with Propofol via an infusion pump (0.22 mg/(kg min)). Analgesia was provided with Dexmedetomidine (0.5 µg/(kg h) to 1 µg/(kg h)) plus Ketamine (0.8 mg/(kg h)). Hydration was maintained with sterile saline (3 mL/(kg h)) and glucose solution (3 mL/(kg h)).

Once anesthetized, the animal’s head was placed in a stereotaxic frame, and the skull was exposed. Stainless steel screw electrodes (1.4 mm diameter) were placed on the surface (over the dura) of different cortical areas (rostral and dorsolateral prefrontal, primary somatosensory, primary auditory, primary motor, primary visual, and parietal posterior). Additionally, depth electrodes were implanted in the medial geniculate nuclei (the auditory thalamus) and the lateral geniculate nuclei (the visual thalamus). The electrodes were connected to a Winchester connector, which, along with two plastic tubes, were attached to the skull with acrylic cement to hold the animals’ heads in a fixed position without pain or pressure.

Once the surgery was completed, the animals were kept in the laboratory to monitor their postoperative progress. After confirming their normal recovery (24 hours), they were transferred to the animal facility, where they received an analgesic every 24 hours for 48 hours (Ketoprofen, 2 mg/kg subcutaneously), or longer if necessary. The incision margins were kept clean, and topical antibiotics were applied daily.

### Experimental protocol

Experimental sessions were conducted between 14 and 18 h in a controlled-temperature environment (21-23 °C). During these sessions (as well as during the adaptation sessions), the animals’ heads were supported in a stereotaxic position by four steel bars placed in the chronically implanted plastic tubes, while the body rested in a sleeping bag (semi-restricted). ECoG and thalamic LFP activity were recorded in a monopolar (referential) configuration, with all channels referenced to a common electrode placed over the left frontal sinus. The electromyogram (EMG) of the nuchal muscles was recorded using a bipolar electrode placed with conductive paste during the experiment. The bioelectrical signals were amplified (×1000), sampled (1024 Hz, 16 bits), and stored on a PC using Spike 2 software (Cambridge Electronic Design).

In the experimental phase, the auditory paradigms were presented on a PC running MATLAB and scripts for their generation. The system’s output was suitable for speakers adapted for cats. The cats were recorded daily for 4 hours, during which the auditory paradigms to be studied were repeatedly presented 6-8 times, with a 900 ms pause between sessions. Two types of cognitive paradigms were used: the Roving paradigm (Canales-Johnson et al., 2021) and the Local-Global paradigm (Bekinschtein et al., 2009). For the latter, we used a variant that omits deviant stimuli (Chennu et al., 2016), although the omission effect was not studied in this work. The experiments of 6-8 sessions were repeated ≈10 to 24 times for each paradigm. The two paradigms were presented on alternating days.

### Stimuli and tasks

#### Local-Global task

The experimental paradigm was implemented following the design proposed by Bekinschtein et al. (2009) and Chennu et al. (2016). In each day the cat was presented 6-9 whole sessions of stimuli consisting of 6 block types, with 10-15 mins rest between sessions. Additionally, each session consisted of 6 block types, with 3 mins rest between blocks. Four of these were experimental blocks, whereas two were control blocks. Audible binaural vowel tones 50 ms long were presented in grouped sequences of 4 or 5 sounds, with short gaps of 100 ms. Individual sounds were vowels sounds “E” and “O”, adapted from (Blume et al., 2022). The stimuli were binaural, and it was presented at an intensity of 60 dB SPL. As shown in Figure 1, in each test the animals heard a series of 5 short sounds: the first 4 were always identical, and the last was identical (EEEEE or OOOOO, local standard) or different (EEEEO or OOOOE, local deviant). In an additional block, instead of a different sound, the omission of the 5th sound (EEEE or OOOO , “omission”) was presented, which also represents a local deviation. The general regularity of each experimental block was defined by the frequency (74% or 13%) of each test type. In this way, 5 conditions can be defined for each test (“local standard and global standard”, “local deviant and global deviant”, “local deviant and global standard”, “local standard and global deviant”, and “omission”). In all five conditions, each experimental block began with between 20 and 30 global standard trials to establish a clear general standard for the animals. An additional block not shown in the figure consists of the omission control condition. In this block, only the omission trial (EEEE or OOOO) is presented throughout the session, so it is not recognized that a sound is missing. In the analysis, omission deviants are contrasted against the omission control. In both conditions, the animal hears the same series of four sounds, but in the deviant condition, it waits for a sound to come, whereas in the control condition, it does not expect to hear one. This attempts to capture the purest (top-down) error signal in the absence of bottom-up.

Approximately 135 sequences were presented in each block, lasting around 3.2 mins. The interval between consecutive sequences was randomly sampled from a uniform distribution of 700 to 1000 milliseconds. The experiment began with a habituation phase consisting of 20 presentations of a recurring 5-sound sequence. This was immediately followed by the test phase, consisting of 115 sequences. Of these, 85 (approx. 74%) were the global standard, and the remaining were rare. There were approximately 15 (13%) of each type of deviant sequence in a block, pseudorandomly interspersed among the global standards. Between 2 and 5 global standards were always presented between each deviant sequence.

Figure 1 illustrates the structure of each deviant sequence, together enabling the well-established local-global paradigm, first introduced by Bekinschtein et al. (2009), to create orthogonal local and global contrasts of predictability across 4 experimental blocks. In this paradigm, global-standard sequences in X blocks are also local standards, whereas global-standard sequences in Y blocks are local deviants. Consequently, the global-standard sequence in an X block becomes the global deviant in the complementary Y block, and vice versa. In this way, we were able to contrast local standards by averaging global standard trials in X blocks and global deviant trials in Y blocks, against local deviants (by averaging global deviant trials in X blocks and global standard trials in Y blocks) to examine the mismatch response.

The dominant vowel type (E or O) of the sequence presented within each block was counterbalanced, resulting in the 4 experimental blocks listed in Table 1. So, for example, in the A-X block (Table 1, first row), the locally standard EEEEE sequence was also the global standard. Rare global deviants in this block were also locally deviant, either the sequence EEEEO, or the omission sequence EEEE , where the fifth tone was omitted. By contrast, in the B-Y block (Table 1, third row), the locally deviant OOOOE sequence was now the global standard. Global deviants in this block were either the locally standard OOOOO sequence or the OOOO omission sequence. The 4 blocks were presented in pseudorandom order.

**Table 1:**
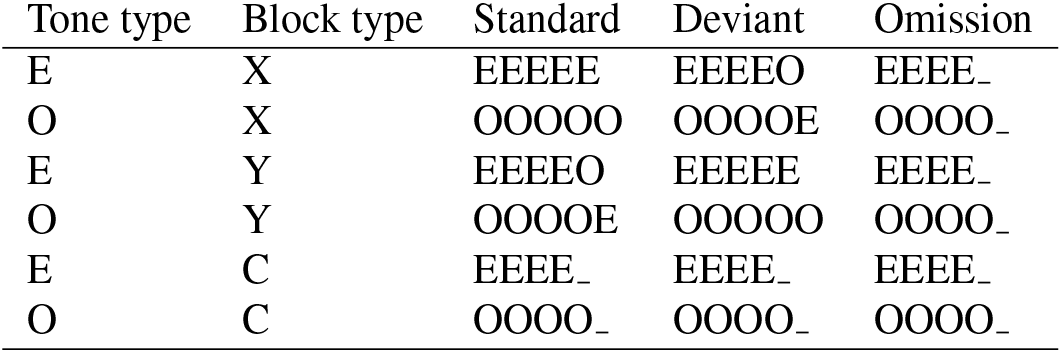
Types of blocks presented in the Local-Global paradigm. The four types of blocks were presented in random order, interspersed with two control blocks.

Two additional control blocks were randomly interspersed among the experimental blocks. These blocks had 120 sequences each, all of which were omitted. Hence, the responses to omissions in the control blocks, referred to as omission controls, served as a baseline for comparison with omissions in the experimental blocks. The two control blocks contained 60 repetitions of the EEEE and OOOO sequences, respectively.

#### Roving task

Repetitive trains of identical tones of varying lengths were presented, drawn from 20 different frequencies ranging from 250 to 6727 Hz in a pseudorandom order. Each tone had a duration of 64 ms, with an interstimulus interval of 503 ms (Figure 1). The stimuli were binaural, ramped pure tones presented at an intensity of 60 dB SPL. Tones were identical within each stimulus train but differed between trains. Consequently, the first tone of each train was considered an unexpected deviant, whereas the final tone was considered an expected standard. In this way, we can explore responses related to the violation regardless of stimulus characteristics (Canales-Johnson et al., 2021).

#### Preprocessing

The data were preprocessed in MATLAB using the EEGLAB toolbox, where they were epoched and band-pass filtered between 0.5 and 200 Hz. For the ERP analysis, the data was filtered between 3 and 30 Hz and notch-filtered at 50, 100, and 150 Hz. The data sets were segmented from -800 to 600 ms relative to the onset of the fifth tone or omission. The segments were then divided into standard epochs and deviant epochs. Epochs containing eye artifacts or movements were removed. Only wakefulness was included in this analysis; therefore, epochs during sleep were removed. The total number of trials per condition per animal ranged from 1076 to 2848. These numbers were achieved by adding trials performed on different days and sessions.

#### Event-related potential (ERP) analyses

After preprocessing, ERPs were computed independently for each recording channel and stimulus condition by averaging epochs time-locked to stimulus onset. Separate averages were obtained for standard and deviant stimuli in the Roving Oddball paradigm and for standard and local deviant trials in the Local Effect paradigm. ERP waveforms from intracranial cortical recordings, thalamic recordings, and scalp electrodes were visually inspected to characterize auditory evoked responses and mismatch activity. The resulting ERPs were subsequently used to quantify the encoding of prediction errors using Mutual Information (MI) and Co-Information (co-I) analyses.

#### Mutual Information (MI) analyses

We employed Gaussian Copula Mutual Information (Ince et al., 2017) to estimate the mutual information between the stimulus class (standard/deviant) and the ECoG response. The GCMI toolbox estimates mutual information after Gaussian copula normalization of the ERP data (for full details of the method (see Ince et al. (2017)), and we have previously implemented it to study neural signals in multiple tasks and species (Äijälä et al., 2026; Blume et al., 2026; Canales-Johnson et al., 2023; Gelens et al., 2024; Olivares et al., 2025; Potash et al., 2025; Roberts et al., 2026). This information-theoretic approach is complemented by non-parametric permutation testing to correct for multiple comparisons. At every time point, the signal was randomly assigned a stimulus class label, and permuted 1000 times for each individual electrode. For each permutation, the maximum MI value across all time points was taken, and the 95th percentile of this value was used as the threshold for significance. With this method, a Familywise Error Rate (FWER) of 0.05 is achieved. Electrodes with significant mutual information between the stimulus class and the ECoG response (BB or ERP) were selected as electrodes of interest for further co-I analyses.

#### Co-Information (Co-I) analyses

Co-information was quantified in three ways: (1) within signals (i.e, single electrodes against themselves), (2) between electrodes within the same cortical area (i.e., between all electrodes within the frontal or temporal cortices), (3) between electrodes in different cortical areas (i,e., between all the temporal and frontal electrodes). Co-information was calculated on a trial-by-trial basis, which results in the quantification of the redundant (positive co-I value) and synergistic (negative co-I value) information that the neural signals encode about the stimulus class.

Co-I is formally expressed as:

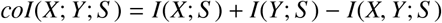

For each time point, *I*(*X*; *S*) corresponds to the mutual information (MI) between the signal at recording site X and stimuli class S. *I*(*Y*; *S*) corresponds to the MI between the signal at recording site Y and stimuli class S. Finally, *I*(*X, Y*; *S*) corresponds to the MI between stimuli class S combining signals from recording sites X and Y. A positive co-information value indicates that signals X and Y contain redundant (i.e., identical) information about the stimulus class S. In contrast, a negative co-information value signifies that signals X and Y contain synergistic information about the stimulus class S. In other words, considering both signals together provides more information than the sum of the information they provide when considered individually (i.e., the overall information provided by the signals is more than the sum of its parts). As with MI, Co-I analyses were complemented with identical non-parametric permutation testing (FWER < 0.05).

#### Multivariate Co-Information (MVCo-I) analyses

To estimate MI and co-I in high-dimensional neural responses, we used Multivariate Co-Information (MVCo-I) (Gelens et al., 2024). MVCo-I combines Multi-Variate Pattern Analysis (MVPA) with information-theoretic co-information to quantify the representational interactions in predictions made from cross-validated multivariate models. In brief, we first apply MVPA in the typical way (here, using the MVPA Light toolbox; (Treder, 2020)) with 10-fold cross-validation (CV). In a 10-fold CV, the overall dataset is randomly separated into 10 disjoint subsets. Then, a model is fit on 9 of those subsets and tested on the 10th, and this is repeated for each of the 10 subsets. Here, we take the decision value of the learned classifier (the linear combination of the weights and the data, which is then thresholded to make the classification) for each test-set trial. This quantifies how strongly the informative pattern the classifier had learned was present in the data on that trial. We combine the test-set decision values across all 10 CV repetitions and compute the mutual information between these out-of-sample decisions and the true stimulus value for each trial (Yan et al., 2023a,b). We use MVPA to reduce multi-channel activity to a single scalar value: the decision value (d-val).

We can repeat the MVPA analysis for each time point of the stimulus-locked epochs. Often, temporal cross-decoding (King and Dehaene, 2014) is employed alongside MVPA to assess the consistency of informative patterns over time. For this method, a classifier is trained at time t, and then tested (in the hold-out test folds) at other times. If it can decode, it shows the same pattern learned at time t, which is information from other times. However, this can only compare between data sets or conditions that are in the same physical space: i.e., we can cross-decode across time within one brain region, but we cannot compare between two different brain regions, because there is no way to apply the linear weights learned in the frontal region to the completely different temporal electrodes. Combining MVPA with co-Information (MVCo-I) overcomes this limitation. We compute co-information between the cross-validated decision values of different classifiers. This admits the same interpretation as for the channel-wise analysis. Redundancy indicates that there is common information accessed by both decoding models. Synergy means a super-additive boost in the information available when considering the pattern activation from both models together. When estimating the joint information for the co-information calculation we take the maximum of the individual region MI (because the data processing inequality tells us this is a lower bound on the information that can be extracted from the joint response), the MI from the combined d-vals (2D signal; this has the advantage of being a low dimensional response for MI calculation, and being the optimal informative signal from each region) and the MI from a joint MVPA model fit to the combination of channels from both regions (1D d-vals, but which has the possibility to include synergistic information between the regions which we want to capture with this measure).

We apply this methodology here in two ways. First, we look at within-area MVCo-I (Figure 4). For this, we train CV classifier models separately at each time point. We then calculate the co-information between two time points using the crossvalidated decision values of the two models. Note that a crucial difference between this and the temporal cross-decoding method is that we always use the learned model to optimally decode information at that time point. Cross-decoding can tell if the same pattern is informative, but we can see redundant information even when the informative pattern changes. We can then compute MVCo-I between regions in the same way (between-areas MVCo-I; Figure 5).

## Supplementary Figures

**Figure S1:**
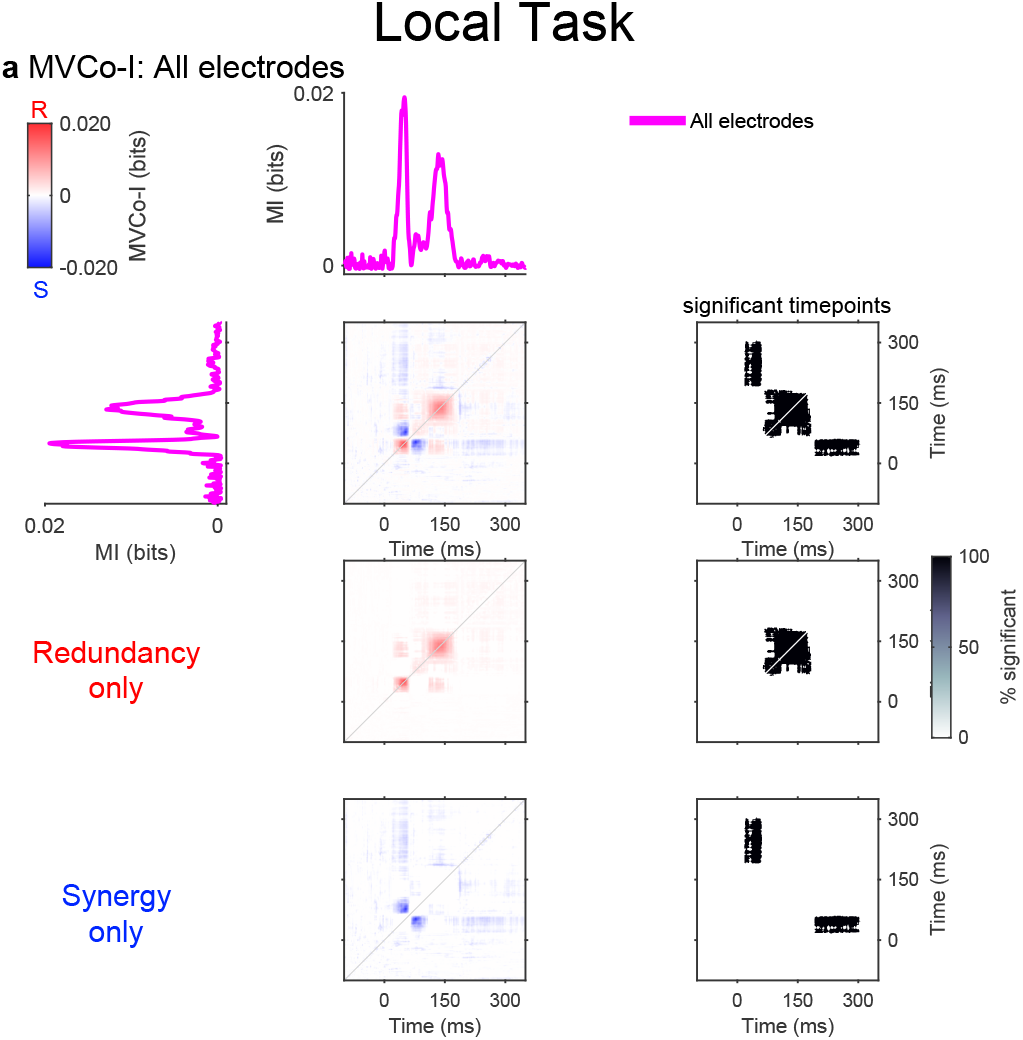
Within-network multivariate redundancy and synergy for cat Mic during the Local Effect of the Global–Local task. All thalamic and cortical electrodes were entered jointly into the multivariate decoder. The magenta trace shows mutual information between the cross-validated decoder decision values and stimulus category. The central matrices show complete MVCo-I, redundancy only, and synergy only from top to bottom. Positive red values indicate redundant information and negative blue values indicate synergistic information. Grey-scale matrices show the binary permutation-test significance mask for this cat. Time is relative to tone onset and spans -100 to 350 ms.

**Figure S2:**
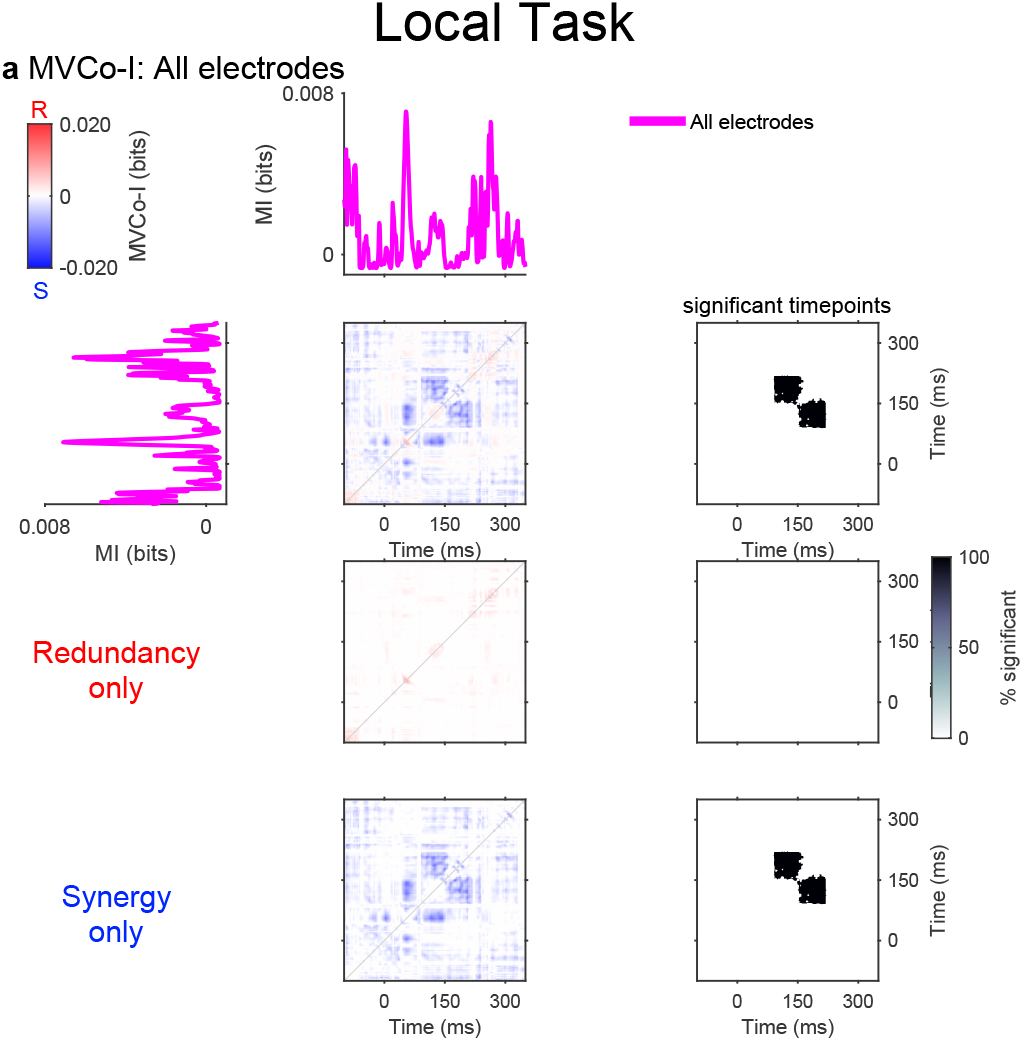
Within-areas multivariate redundancy and synergy for cat Neg during the Local Effect of the Local-Global task. All thalamic and cortical electrodes were entered jointly into the multivariate decoder. The magenta trace shows mutual information between the cross-validated decoder decision values and stimulus category. The central matrices show complete MVCo-I, redundancy only, and synergy only from top to bottom. Positive red values indicate redundant information and negative blue values indicate synergistic information. Grey-scale matrices show the binary permutation-test significance mask for this cat. Time is relative to tone onset and spans -100 to 350 ms.

**Figure S3:**
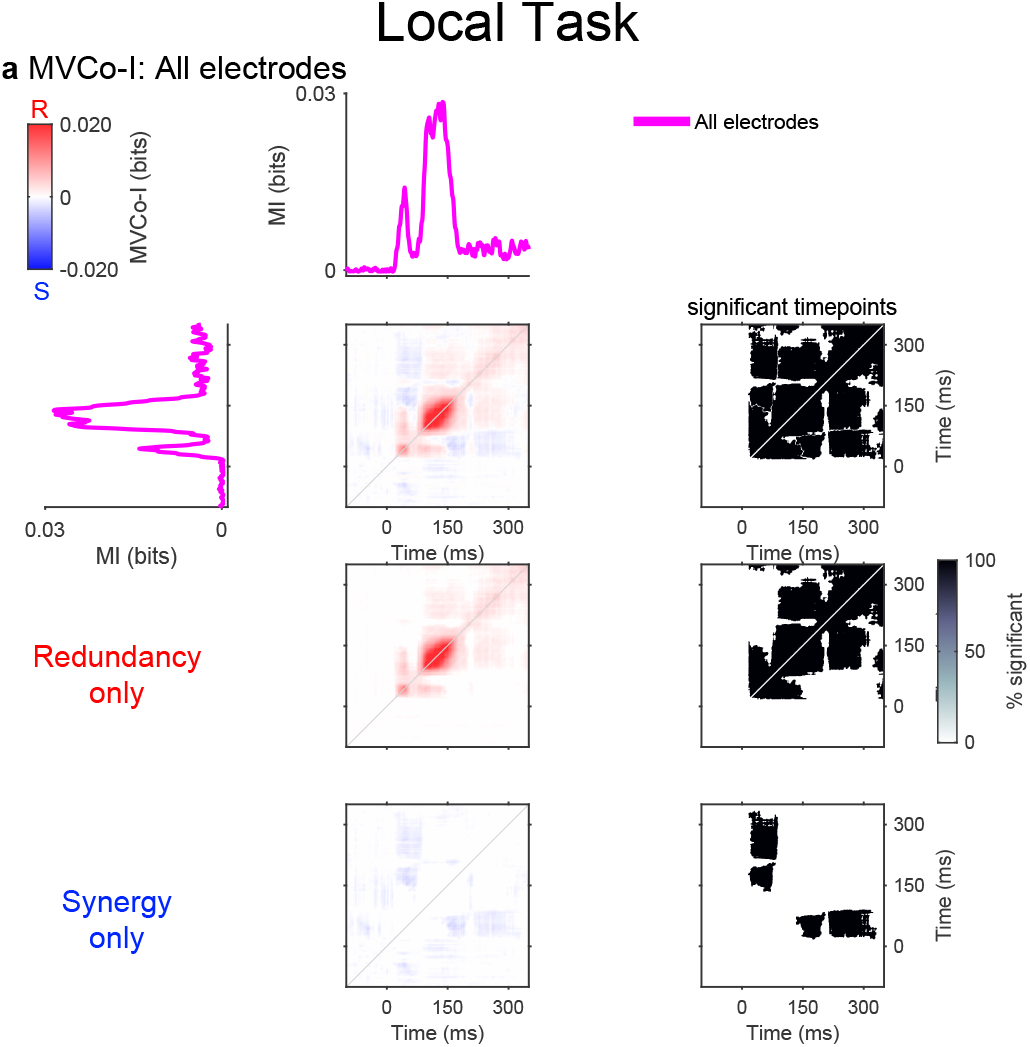
Withi-areas multivariate redundancy and synergy for cat Nim during the Local Effect of the Local-Global task. All thalamic and cortical electrodes were entered jointly into the multivariate decoder. The magenta trace shows mutual information between the cross-validated decoder decision values and stimulus category. The central matrices show complete MVCo-I, redundancy only, and synergy only from top to bottom. Positive red values indicate redundant information and negative blue values indicate synergistic information. Grey-scale matrices show the binary permutation-test significance mask for this cat. Time is relative to tone onset and spans -100 to 350 ms.

**Figure S4:**
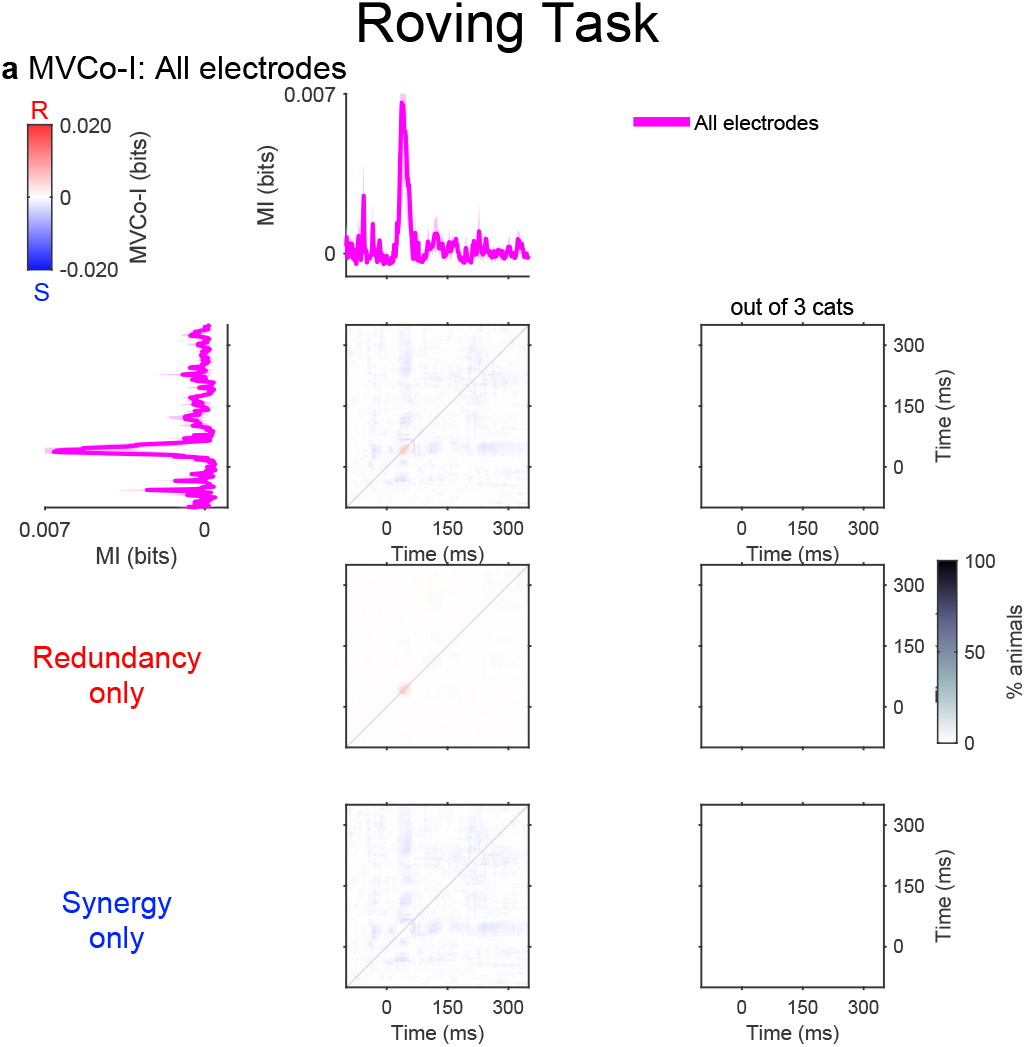
Within-areas multivariate redundancy and synergy averaged across all three cats during the Roving Oddball task. All thalamic and cortical electrodes were entered jointly into the multivariate decoder. The magenta trace shows mutual information between the cross-validated decoder decision values and stimulus category. Shading around the MI trace indicates SEM. The central matrices show complete MVCo-I, redundancy only, and synergy only from top to bottom. Positive red values indicate redundant information and negative blue values indicate synergistic information. Grey-scale matrices show the proportion of cats with significant MVCo-I at each time-point combination. Time is relative to tone onset and spans -100 to 350 ms.

**Figure S5:**
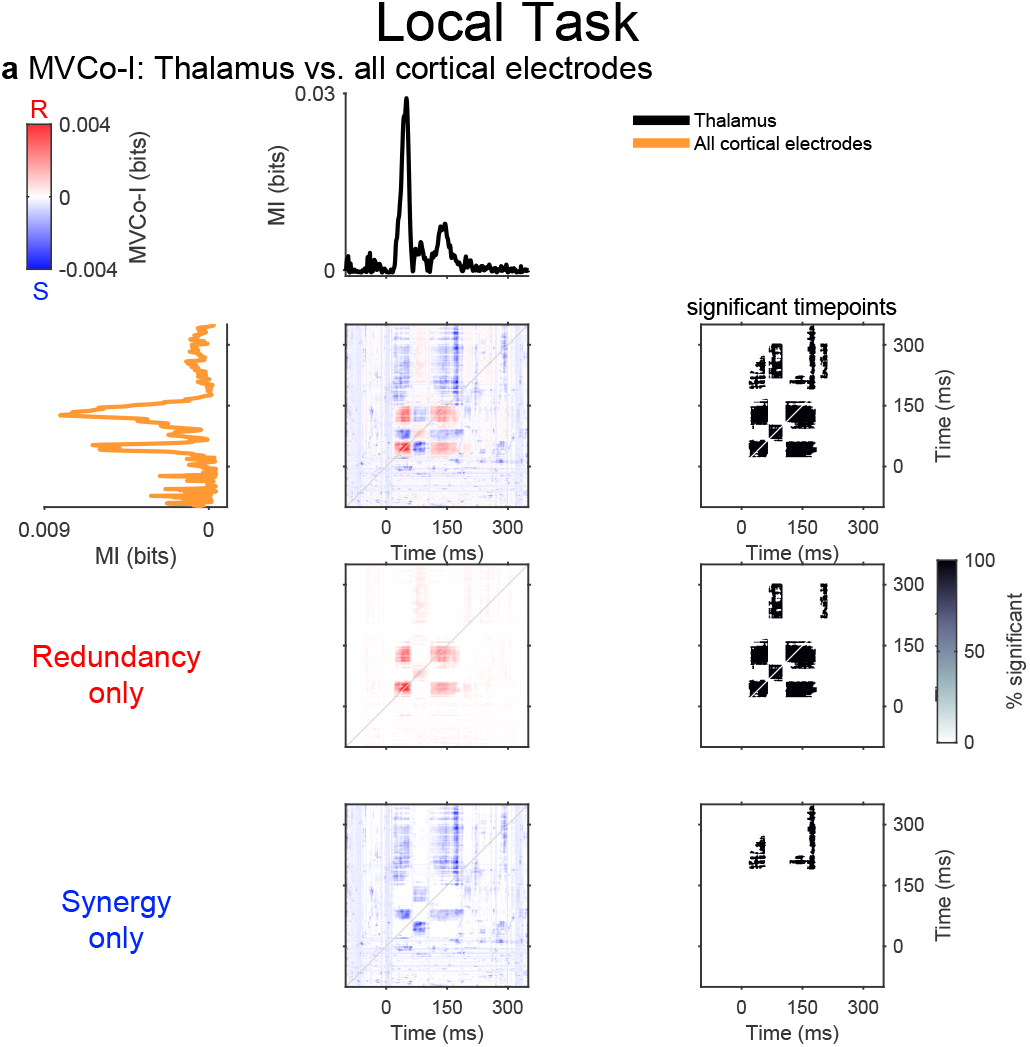
Multivariate redundancy and synergy between thalamic and cortical prediction-error representations for cat Mic during the Local Effect of the Local-Global task. Separate multivariate decoders used thalamic electrodes and all cortical electrodes. Black and orange traces show thalamic and cortical mutual information, respectively. The central matrices show complete cross-regional MVCo-I, redundancy only, and synergy only from top to bottom. Positive red values indicate redundant information and negative blue values indicate synergistic information. Grey-scale matrices show the binary permutation-test significance mask for this cat. Time is relative to tone onset and spans -100 to 350 ms.

**Figure S6:**
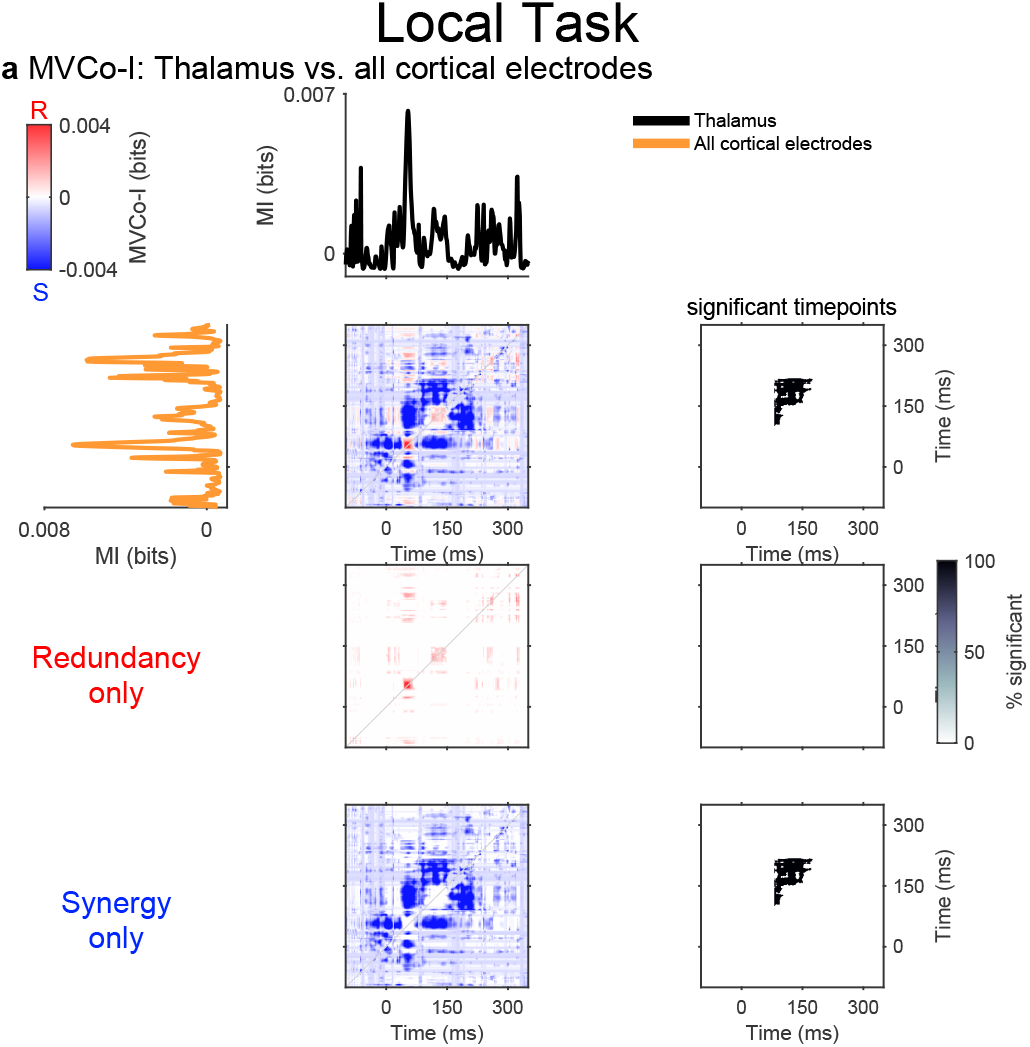
Multivariate redundancy and synergy between thalamic and cortical prediction-error representations for cat Neg during the Local Effect of the LocalGlobal task. Separate multivariate decoders used thalamic electrodes and all cortical electrodes. Black and orange traces show thalamic and cortical mutual information, respectively. The central matrices show complete cross-regional MVCo-I, redundancy only, and synergy only from top to bottom. Positive red values indicate redundant information and negative blue values indicate synergistic information. Grey-scale matrices show the binary permutation-test significance mask for this cat. Time is relative to tone onset and spans -100 to 350 ms.

**Figure S7:**
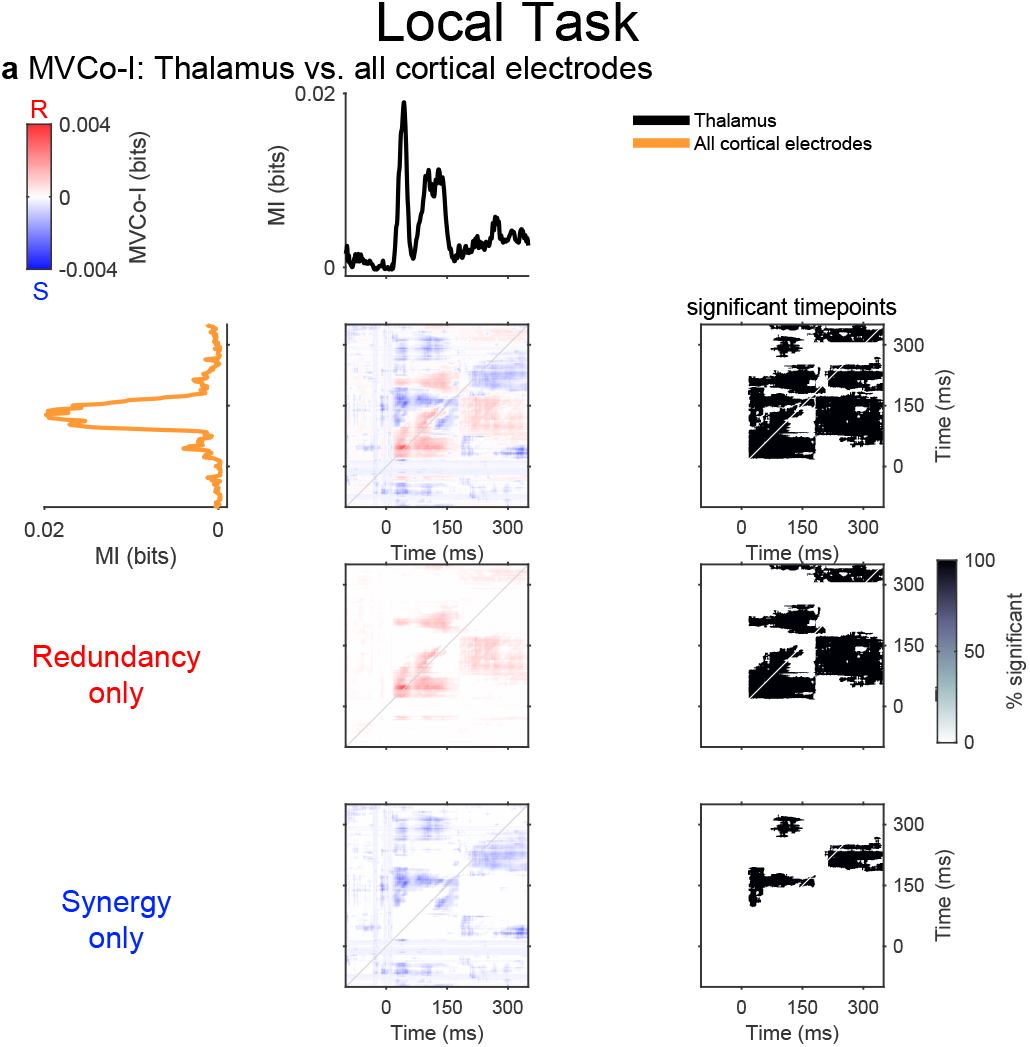
Multivariate redundancy and synergy between thalamic and cortical prediction-error representations for cat Nim during the Local Effect of the Local-Global task. Separate multivariate decoders used thalamic electrodes and all cortical electrodes. Black and orange traces show thalamic and cortical mutual information, respectively. The central matrices show complete cross-regional MVCo-I, redundancy only, and synergy only from top to bottom. Positive red values indicate redundant information and negative blue values indicate synergistic information. Grey-scale matrices show the binary permutation-test significance mask for this cat. Time is relative to tone onset and spans -100 to 350 ms.

**Figure S8:**
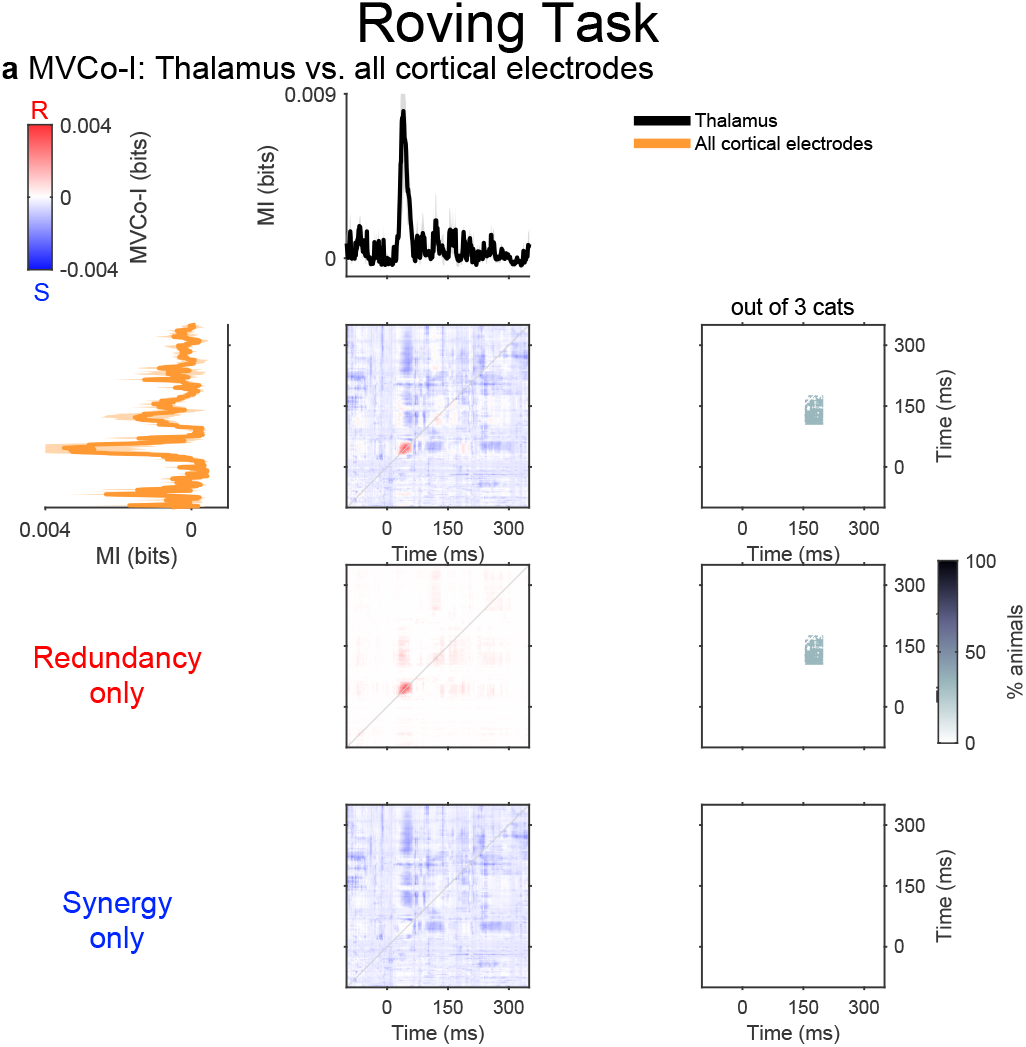
Multivariate redundancy and synergy between thalamic and cortical prediction-error representations averaged across all three cats during the Roving Oddball task. Separate multivariate decoders used thalamic electrodes and all cortical electrodes. Black and orange traces show thalamic and cortical mutual information, respectively. Shading around the MI traces indicates SEM. The central matrices show complete cross-regional MVCo-I, redundancy only, and synergy only from top to bottom. Positive red values indicate redundant information and negative blue values indicate synergistic information. Grey-scale matrices show the proportion of cats with significant cross-regional MVCo-I. Time is relative to tone onset and spans -100 to 350 ms.

